# Structural consequences of deproteinating the 50S ribosome

**DOI:** 10.1101/2022.03.10.483802

**Authors:** Daniel S. D. Larsson, Kanchugal P Sandesh, Maria Selmer

**Affiliations:** Department of Cell and Molecular Biology, Uppsala University, Box 596, SE 751 24 Uppsala, Sweden

**Author notes:** Address correspondence to Maria Selmer,. MAX IV Laboratory, Fotongatan 2, SE 224 84 Lund, Sweden.

## Abstract

Ribosomes are complex ribonucleoprotein particles. Purified 50S ribosomes subjected to high-salt wash, removing a subset of ribosomal proteins (r-proteins), were early shown competent for *in vitro* assembly into functional 50S subunits. We here used cryo-EM to determine the structure of such LiCl core particles derived from *E. coli* 50S subunits. A wide range of complexes with large variation in extent of ordered 23S rRNA and occupancy of r-proteins could be identified, and resolved to between 2.8 Å and 9 Å resolution. Many of these particles showed high similarity to *in vivo* and *in vitro* assembly intermediates, supporting the inherent stability or metastability of these states. Similar to states in early ribosome assembly, the main class showed ordered density for 23S rRNA domains 0, I, II, III, VI and the 5’-half of domain IV. In addition, smaller core particles were discovered, which show that the most stable part of the 50S under high-salt conditions includes parts of domain 0 and most of domains I, III and the 5’-half of domain IV and four to eight r-proteins. Our data support a multi-pathway disassembly process based on independent folding blocks, similar but reverse to the assembly process. The study provides examples of dependencies between complex tertiary RNA structure and RNA-protein interactions where protein extensions dissociate before the globular domains. We observe formation of a non-native RNA structure upon protein dissociation, demonstrating that r-proteins stabilize native RNA structure and prevent non-native interactions also after folding.

**IMPORTANCE:** Ribosome assembly and stability remain only partially understood. Incubation of ribosomes with salts was early shown to induce dissociation of the more loosely bound ribosomal proteins (r-proteins) and formation of so-called core particles. In this work, cryo-EM imaging of 50S LiCl core particles from *E. coli* for the first time allowed structural characterization of such particles of different size. The smallest particles demonstrate what constitutes the smallest stable core of the 50S ribosomal subunit, and the sequential comparison with larger particles show how the ribosome disassembles and assembles in layers of rRNA structure stabilized by globular domains and extended tails of r-proteins. Major insights are that ribosomes disassemble along different paths, that dissociation of r-proteins can induce misfolding of rRNA and that extended tails of r-proteins dissociate from rRNA before the globular domains. The characterized particles can be used in future mechanistic studies of ribosome biogenesis.

## INTRODUCTION

The ribosome is a large macromolecular complex translating mRNA into protein in all domains of life. Its intricate assembly is in all organisms a tightly regulated process that has been fine-tuned by evolution. In *E. coli*, 54 ribosomal proteins (r-proteins) and three ribosomal RNAs (rRNAs) fold and assemble with the help of a multitude of auxiliary RNA modification enzymes and ribosome assembly factors (reviewed in (1, 2)). Despite the complexity of the process, the assembly time for a mature ribosome *in vivo* in exponentially dividing *E. coli* has been estimated to only 2 minutes (3).

The structure and assembly of the 50S subunit has been extensively studied *in vitro*, and methods were early developed to reconstitute active subunits from components (4). Another *in vitro* approach was to promote the dissociation of r-proteins through high-salt wash of ribosomal particles. Proteins were shown to dissociate from the core at defined concentrations of *e*.*g*. LiCl (4, 5). The observed order of leaving r-proteins nearly agrees with the reverse of the *in vivo* assembly order followed by quantitative mass spectrometry (6) as well as the *in vitro* reconstitution protocol (7). A subset of the of the ribosomal proteins that dissociate during salt wash also exchange *in vivo*, presumably as a mechanism by which damaged proteins can be continuously replaced (8).

Our structural understanding of folding and assembly of ribosomes has greatly benefitted from progress in cryogenic electron microscopy (cryo-EM), where classification schemes in single-particle reconstruction have proven essential to study these heterogeneous particle ensembles. To enrich *in vivo* ribosome assembly intermediates for these studies, bacteria have been subjected to assembly inhibitors (9, 10), knock-down of r-proteins or assembly factors (11–14) and pull-downs with assembly factors as bait (15, 16).

In one such study, a bL17 depletion strain was used to generate a multitude of ribosomal 50S sub-particles (11), allowing identification of several cooperative folding blocks. In agreement with previous studies (10), a model of three major routes of LSU folding upon bL17 perturbation was proposed, where the central protuberance (CP), the particle base and the L7/L12 stalk formed in different order.

Structural studies of assembly intermediates from LSU *in vitro* reconstitution (17) showed five distinct precursors that shared many features with *in vivo* assembly intermediates from the bL17-depleted strain. The later intermediates displayed an intricate variety of folding states of the protuberances and the peptidyl transfer center (PTC).

Salt-washed LiCl core particles have been demonstrated to be assembly-competent, allowing reconstitution into active 50S subunits (4, 7). They also work as *in vitro* substrate for RlmF, an early rRNA modifying enzyme that lacks activity on naked RNA (18), supporting at least local similarity to states occurring during ribosome assembly *in vivo*. There is to date no high-resolution structure of these particles, but early negative-stain electron microscopy showed high degree of heterogeneity where approximately 40% of the ribosomes remained as compact particles lacking the three characteristic protuberances of the LSU (19).

We here set out to further characterize the structure of LiCl core particles using single-particle cryo-EM, with the goal to at high-resolution characterize the structure and composition of these particles. We identified 6 major types of 50S sub-particles with a wide range of sizes–and a multitude of variants of each of these. The method allowed observation of core particles smaller than so far identified 50S assembly intermediates and the opportunity to examine effects of removing particular r-proteins. Comparison of these particles enabled us to identify stability dependencies between rRNA secondary structure elements and r-proteins, to reconstruct possible disassembly and assembly pathways and to demonstrate strong similarities between intermediates formed during disassembly and assembly.

## MATERIAL AND METHODS

### Preparation and characterization of LiCl core particles

Ribosomes were purified from strain JW5107 (*ΔybiN*) of the Keio collection (20). Tight-coupled 70S ribosomes were purified according to (21) with minor modifications and separated into subunits. 50S subunits were subjected to a high-salt wash in 3.5 M LiCl and the resulting core particles purified on a sucrose gradient (details in Supplementary methods). Methylation activity of RlmF on LiCl core particles was determined using a tritium labelling assay (details in Supplementary methods).

### Cryo-EM

In summary, 3 uL sample with 160 nM ribosome concentration was incubated for 30 seconds on grids with holey carbon foil covered with 2-nm continuous carbon and blotted for 3.5 seconds before plunge frozen in liquid ethane. Data was collected at 300 kV and 75,000x magnification with 0°, 15° and 30° sample tilt with a total dose of 41.0 or 42.2 e^-^/Å^2^. Data was processed using RELION (22, 23). Detailed procedure in Results and Supplementary methods.

### Structure analysis

PDB model 4YBB (24) was refined into the highest-resolution map. Flexible or missing parts were removed from the model. Occupancy was estimated by calculating the average map value at the atomic locations of r-proteins and rRNA helices, and by manual inspection. Clustering of correlated structural elements was calculated using an in-house developed script; see Supplementary methods for details.

### Data Availability

Atomic coordinates for the consensus structure have been deposited in the Protein Data Bank under accession code <u>7ODE</u>. All maps have been deposited in the Electron Microscopy Data Bank with accession codes <u>EMD-12826</u> for the consensus reconstruction and <u>EMD-12828</u> to <u>EMD-12841</u>, <u>EMD-12843</u> and <u>EMD-12844</u> for the different classes.

## RESULTS

### Sample preparation

LiCl core particles were prepared from active 50S subunits by incubation with 3.5 M LiCl buffer, purified on a sucrose gradient (Figure S1A) and enriched for RlmF substrate by pull-down with His-tagged A1618 rRNA methyltransferase RlmF (Figure S1B). LiCl core particles could be methylated *in vitro* by RlmF, as previously shown (18). Using tritium-labelled S-adenosyl methionine, methylation showed saturation after 10 minutes at ∼20% modification (Figure S2). RlmF was by SDS-PAGE judged to be present at approximately the same stoichiometry as uL2 (Figure S1C) and LC-MS/MS analysis showed similar or higher signal as for the stably bound r-proteins. However, no distinct density for the protein could be found in any of the classes.

### High-resolution cryo-EM reconstruction of the 50S LiCl core particle

Cryo-EM imaging, single-particle reconstruction and a multitiered classification scheme was used to determine the structure of the LiCl core particle (Figure 1). Because of strong preferred orientation (19), data collected at 15° and 30° tilt were used to improve the angular sampling. The micrographs showed heterogenous particles (Figure S3A,D) and several of the 2-D class averages had one disordered side (Figure S3G). To avoid bias (25), particle-picking was done without templates and the starting 3D model was generated *ab initio*.

**Figure 1.**
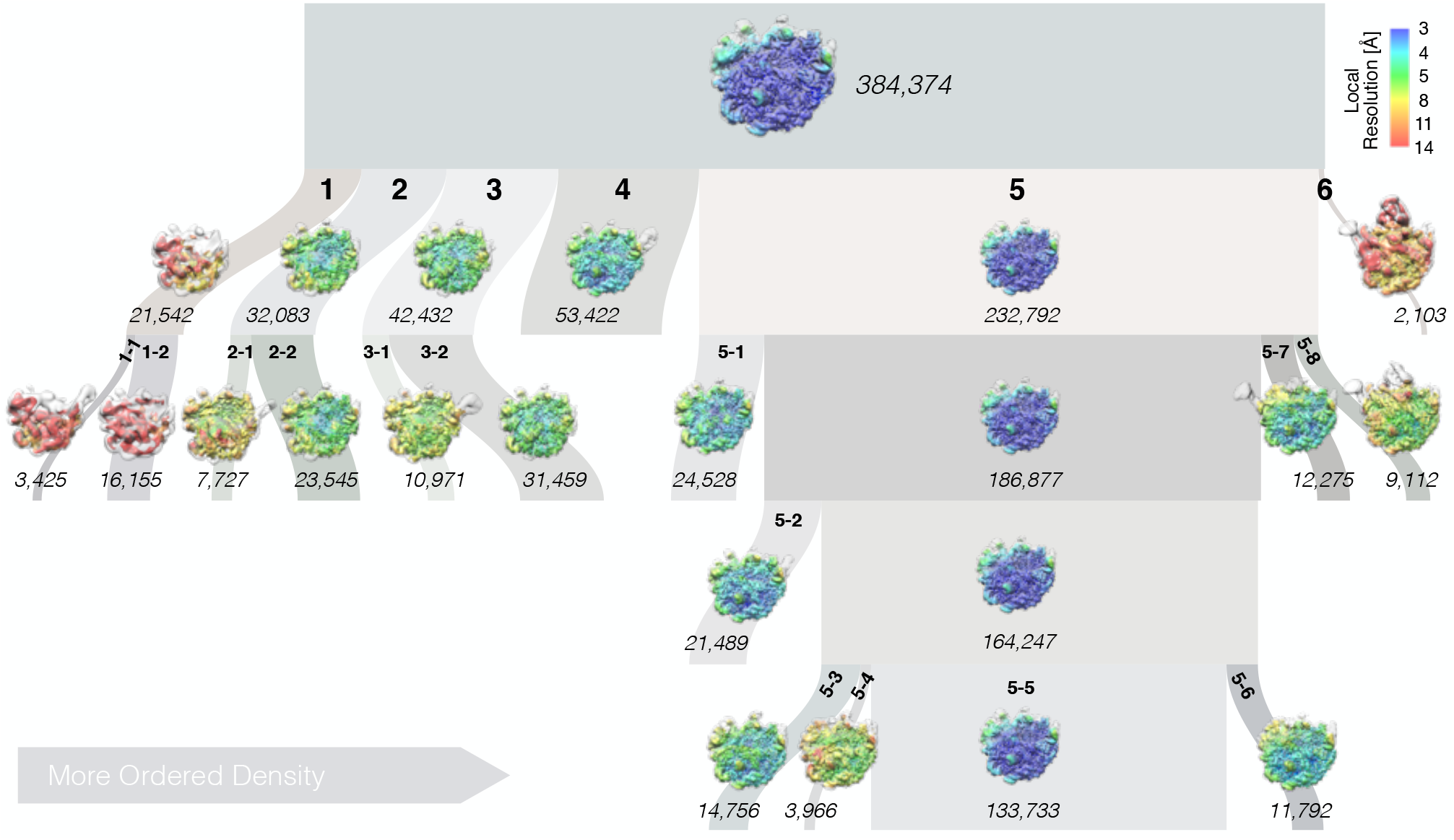
Single-particle reconstruction and hierarchical classification of LiCl core particles. The width of the branches in the diagram reflects the number of particle images (italic font) in each class (bold font) in the tiered classification scheme. Classes are sorted from left to right approximately according to the amount of ordered density. Images show the reconstructions in “crown view” with the L1 protuberance to the left and the L7/L12 stalk to the right (*cf*. Figure S5 for details). Minor classes that did not produce a stable reconstruction by themselves, and hence were discarded, are not shown.

### LiCl washing of the LSU produces a range of sub-particles

A consensus reconstruction from ∼384,000 particle images produced a map at 2.84 Å resolution (Figures 2, S4). The reconstructed density is dome-like with the ordered part at the 50S solvent side, centered on the expanded nascent-chain exit tunnel (Figures 2H-I, 3A,C). None of the three characteristic protuberances of the LSU are ordered.

**Figure 2.**
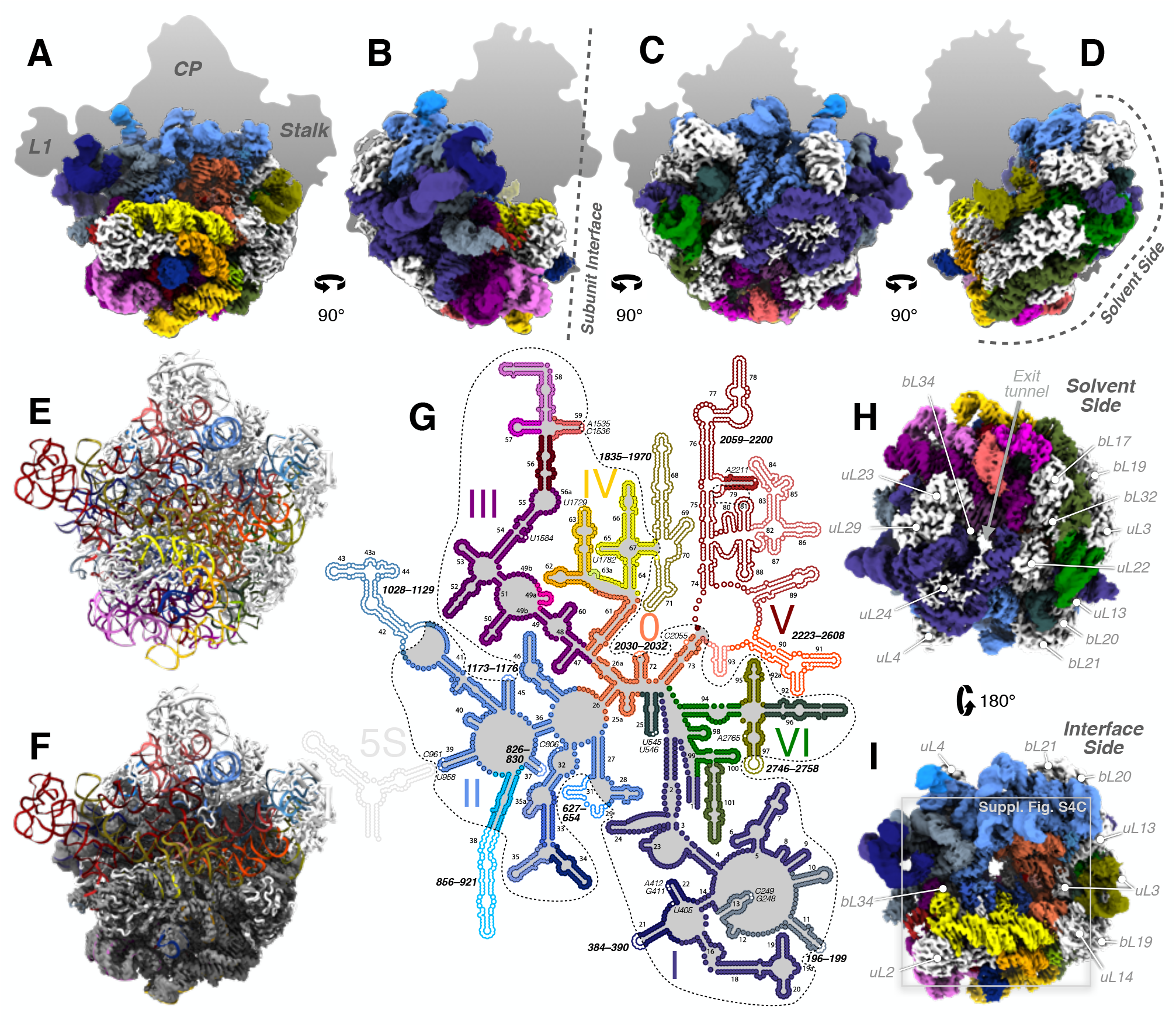
Consensus reconstruction of the 50S LiCl core particle (**A)** The map in “crown view” with the rRNA colored according to (G) and r-proteins in white. The silhouette shows the extent of the mature 50S particle. (**B-D**) Same as (A), rotated in 90° increments around the Y-axis. (**E**) Model of the mature 50S particle (PDB ID 4YBB) in the same view as in (A). RNA colored as in (G) and 5S rRNA and r-proteins in white. (**F**) Model in (E) rigid-body docked in map (gray surface). (**G**) Secondary-structure map of the LSU rRNA (modified from (37)). Folded parts in the consensus particle are indicated with filled circles and outlined with a dashed line. Disordered helices, loops or single nucleotides are shown as white dots and labeled with italic script. The SSE elements are colored in orange, indigo, blue, purple, yellow or red tones for domains 0–VI, respectively. (**H**,**I**) Same as (A) but rotated around the X-axis and viewed along the expanded nascent chain exit tunnel, from (**H**) the solvent-side and (**I**) the interface side. Density for r-proteins (white) indicated. The map is locally lowpass filtered according to the local resolution and displayed at level 0.1. Square indicates the field-of-view of Figure S4C.

Classification resulted in six major classes, numbered 1–6 according to size (Figures 1, S5). Upon further classification, major classes 1–3 split in two, and class 5, the most populated major class, spawned an extensive tree of sub-classes. After four levels of classification, the largest sub-class 5-5 collected approximately 1/3 of the initial particles. Model refinement (Figures 3A–B, S4D) and some of the analysis and figures are based on the consensus map, which is very similar to class 5-5; importantly all conclusions are also valid for this particle.

**Figure 3.**
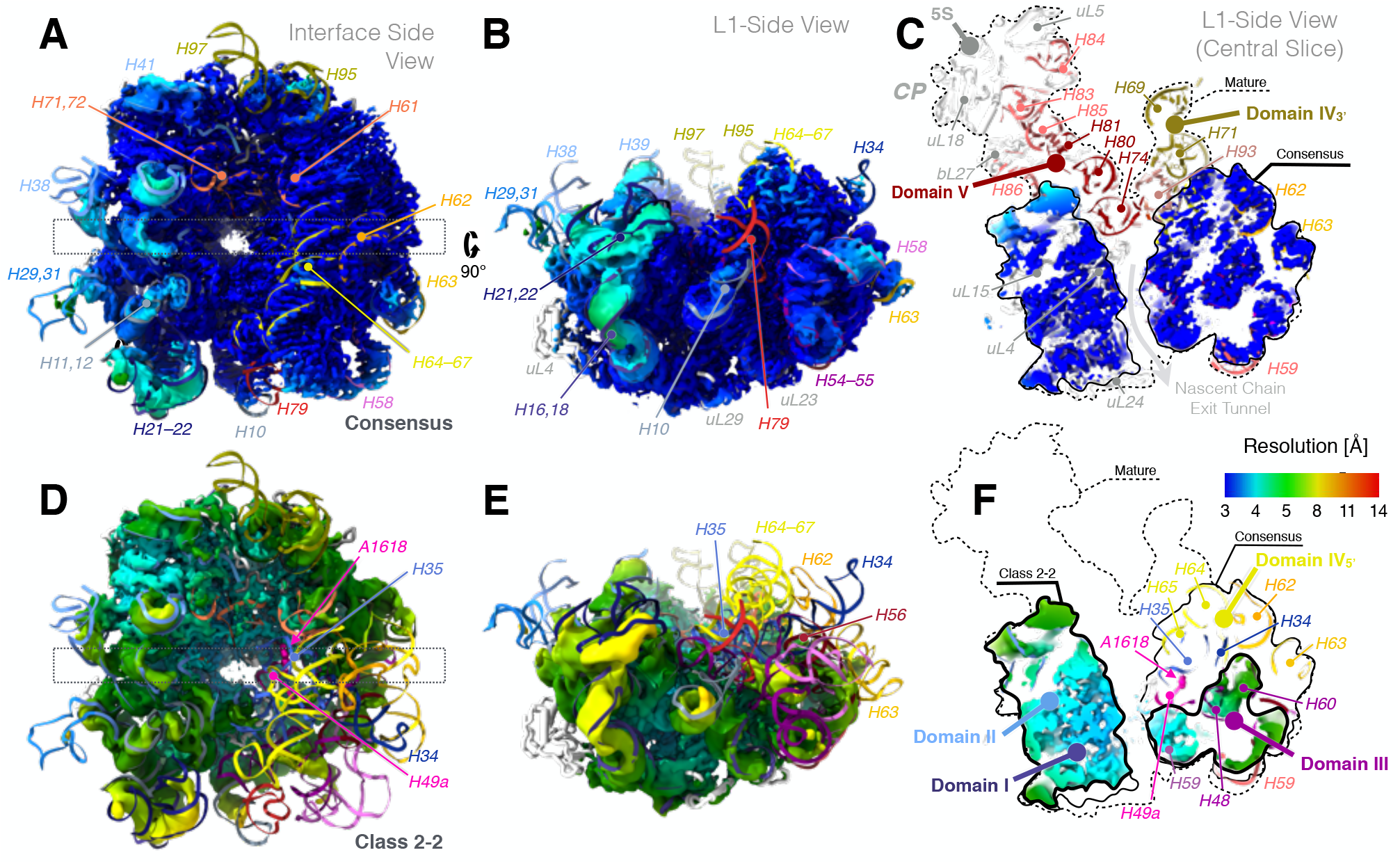
Local resolution and the expanded exit tunnel. (**A**) Consensus reconstruction seen from the interface-side, centered on the nascent-chain exit tunnel, colored according to local resolution, with the refined atomic model colored according to Figure 2G. (**B**) As (A), rotated 90°. (**C**) Central slice through the map (rectangle in (A)) in the same orientation as in (B) showing the expanded exit-tunnel and the absent CP. The rigid-body fitted model of the mature 50S particle (PDB ID 4YBB) is colored as in Figure 2G. The solid black line indicates the extent of the ordered density and the dashed line indicates the extent of the mature particle. (**D-F**) Class 2-2 in same views as in (A-C). The model is the same as in (A-B). In this particle, sub-domain IV_5’_ and parts of domains II, III are disordered compared to the consensus structure, in particular nucleotide A1618 in H49a. The thick black line in (F) indicates the extent of the density in class 2-2, while the thin and the dashed lines are the same as in (C). All maps are locally lowpass filtered according to the local resolution and are displayed at level 0.13.

Most classes could be resolved to better than 4.5 Å resolution, but the smallest and largest particles could only be resolved to approximately 9 Å resolution (Figures S6–S8).

### The LSU is the most stable at the solvent-side

In the majority of the particles, more than half of the 23S rRNA is folded, including domains 0, I, III, VI, most of domain II and the 5’ half of domain IV (*i*.*e*. H62–H67, from now on referred to as sub-domain IV_5’_) (Figure 2G). The folded regions are at the solvent-side and particle base of the LSU and to a high degree correlate with the presence of r-proteins (Figures 2H–I, S4B). Several rRNA helices appear as folded but flexible low-resolution features (Figure 3A–B). The three major protuberances and the subunit interface, including functionally important regions such as the PTC and binding sites for tRNAs and GTPases, are unstructured (Figure 2). In addition, many single nucleotides involved in native tertiary contacts are disordered (Figure 2G).

### The LiCl core particle classes are analogous to *in vivo* and *in vitro* assembly intermediates

Major class 1 is the so far smallest identified stable core of the LSU. Its low resolution is probably due to structural flexibility and heterogeneous presence of r-proteins. Classes 2 through 6 show strong similarities to *in vitro* reconstitution intermediates (17) and to *in vivo* assembly intermediates under bL17 depletion (11) (Figure S9), which is remarkable since they all include bL17. In line with previous structures, the particle base is stable in absence of folded protuberances.

To group the particle classes and to identify cooperativity between binding of r-proteins and folding of rRNA, automated (Figures S10, S11) and manual (Table S1) occupancy estimations were subjected to hierarchical clustering (Figure S12A–D). The classes grouped into smaller (classes 1, 2, 3 and 5-1), medium-sized (classes 4 and 5, except 5-1 and 5-8) and larger particles (classes 5-8 and 6) (Figure S12I), similar to early, intermediate and late 50S assembly intermediates (Figure S9). In the small particles, most of domains 0, I, III, VI and a core of domain II is folded. In the medium-sized particles, more of domain II, as well as sub-domain IV_5’_ and parts of domain V are folded. The large particles are mature-like, but parts of sub-domain IV_3’_ and parts of domain V are unfolded. Clustering of structural features (rows in Figure S12A–D) approximately agrees with previously identified folding blocks (11).

### The misfolded 3’ strand of helix H73

Non-native density at the base of the L7/L12 stalk shows up in several classes (1-1, 2-1, 3-1, 4 and 5-2), as a low-resolution protruding loop at low map thresholds (Figures 4, S5, star). One arm of the loop consists of H97 (Figure 4D), slightly shifted compared to the native particle. The other arm, with appearance of an rRNA stem-loop, emerges outside the particle between H1, H94 and H97. Intriguingly, H73 is natively positioned just inside in the “non-star” particles, but absent in all of the “star” classes (Figure S13). Helix H73 is at a 4-way junction, but the region of its 3’ strand (Figure 4B) can, according to secondary structure prediction, form an alternative stem loop (Figure 4C). A predicted 3D model of the stem loop is compatible in size with the observed non-native density (Figure 4D). Closing the loop, an unidentified protein seems to bind to the top of helices H97 and the misfolded H73. Possibly, this could be uL6, which natively binds H97 and is detected at low levels in the LC-MS/MS data (Table S1).

**Figure 4.**
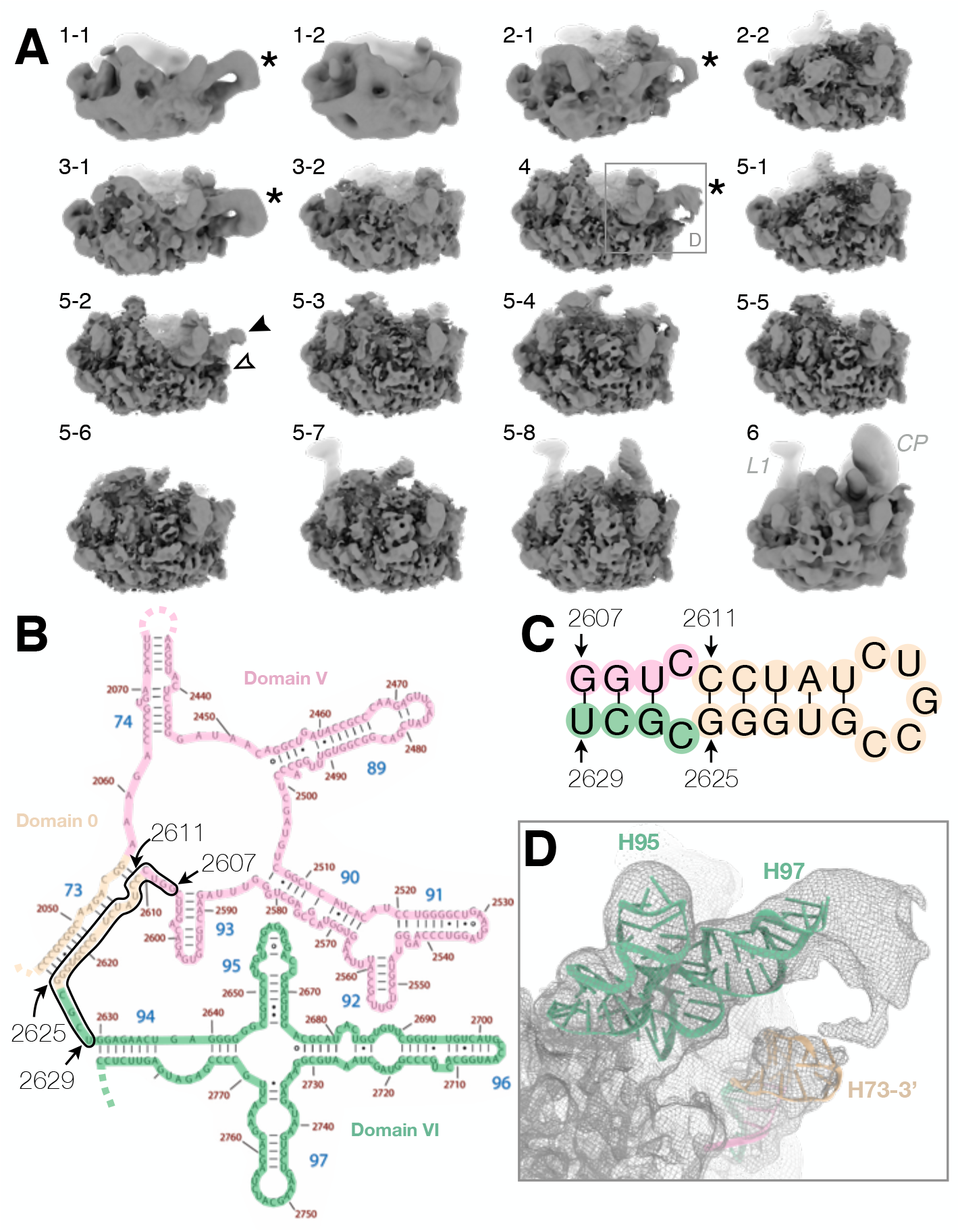
Non-native density close to the base of the L7/L12 stalk. (**A**) In classes 1-1, 2-1, 3-1 and 4, the density of H97 extends and loops back close to H1 (asterisk) at lower map thresholds. In class 5-2, density for the upper (black arrow-head) and lower (white arrow-head) arm are not connected. Analogous classes 1-2, 2-2, 3-2, 5-1 and 5-3, respectively, do not show this extra density. Maps are filtered according to the local resolution and are displayed at level 0.05. (**B**) Secondary structure map of the *E. coli* 23S rRNA shown for parts of domains 0, V and VI (modified from (37)). Omitted segments of the secondary structure map are indicated with dotted lines. A black line encloses the segment that were used for secondary-structure prediction in (C). (**C**) Secondary-structure of the 3’-strand of H73 and neighboring nucleotides predicted by RNAfold. Stem-loops with a 5-base loop and either an 8-bp or a 5-bp stem. Colors as in (B). (**D**) Loop density for class 4 (rectangle in (A)) with a rigid-body docked model of the 8-bp stem loop in (C) predicted by 3dRNA (38) colored as in (C). Models of H97 and H95 are shown for reference.

### R-proteins leave the 50S according to *in vivo* assembly groups

A number of proteins appears to be present across all classes (uL3, uL4, bL17, bL20, bL21, uL22, uL23, uL29 and bL34) and in most cases also uL13 and uL24 (Table S1, Figure S14). All of these except bL32 have firm support in LC-MS/MS data (Table S1). These proteins are all early binders *in vivo*, as identified by pulse-labeling coupled with MS analysis (6) (Figure S14). Furthermore, four proteins that are only found in larger particles (uL2, uL14, bL19 and bL32) bind in the subsequent step *in vivo*.

None of the classes showed clear density for uL1, uL6, bL9, uL10, uL11, bL12, uL15, uL16, bL25, bL27, bL28, bL31, bL33, bL35 or bL36 – although diffuse density prevented unambiguous assessment of some proteins. Trace levels of uL1, uL5, uL6, bL9, uL11, uL15 and bL31 were detected by LC-MS/MS (Table S1), suggesting that some of these r-proteins remain bound to disordered rRNA.

The high solubility of the CP-associated r-proteins uL5, uL18 and bL25 destabilizes the CP in the LiCl washed ribosomes. The CP and the 5S rRNA are only discernable in the largest particle classes, 6 and 5-8 (Figure S5) together with uL5 and uL18 at low occupancy in class 6, and uL18 in class 5-8.

For the three largest r-proteins in the consensus reconstruction (uL2, uL3 and uL4), the long loops and tails that in the native particle interact with the 23S rRNA are invisible (Figure S4C). The interacting rRNA helices are in some cases unstructured and in other cases folded, but in a non-native position.

### Comparison of particle classes show dependencies between different structural elements

Structural consequences of removal of certain r-proteins and of unfolding of rRNA was analyzed by pair-wise comparison between the particle classes, based on the clustering analysis. This revealed folding and stability dependencies that are mostly additive (Figure 5, 6).

**Figure 5.**
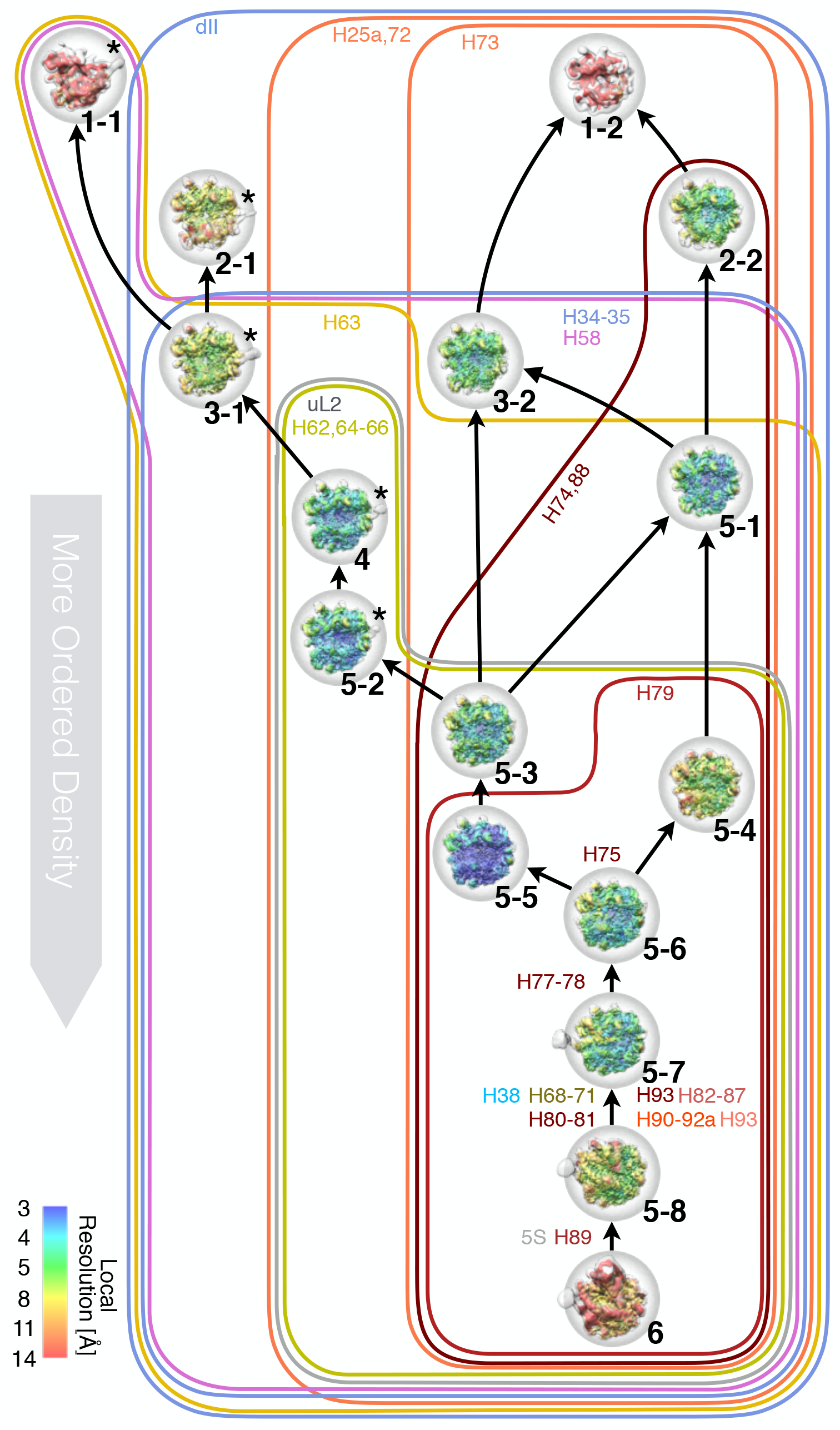
Relationship between observed particle classes based on presence of structural features. Arrows indicate decreasing order. Lines (colored as in Figure 2G) enclose particles with significant density for specific rRNA helices or r-proteins; labels indicate rRNA helices that unfold or r-proteins that dissociate. Particles are shown as in Figure S5. The left branch (5-2, 4, 3-1, 2-1, 1-1) consists of particles with misfolded H73 (asterisks, Figure 4). The right side contains one path with stronger density for uL2, H34–35 and sub-domain IV_5’_ (5-5, 5-3, 3-2) and one path with stronger density for domain V (5-4, 5-1, 2-2).

**Figure 6.**
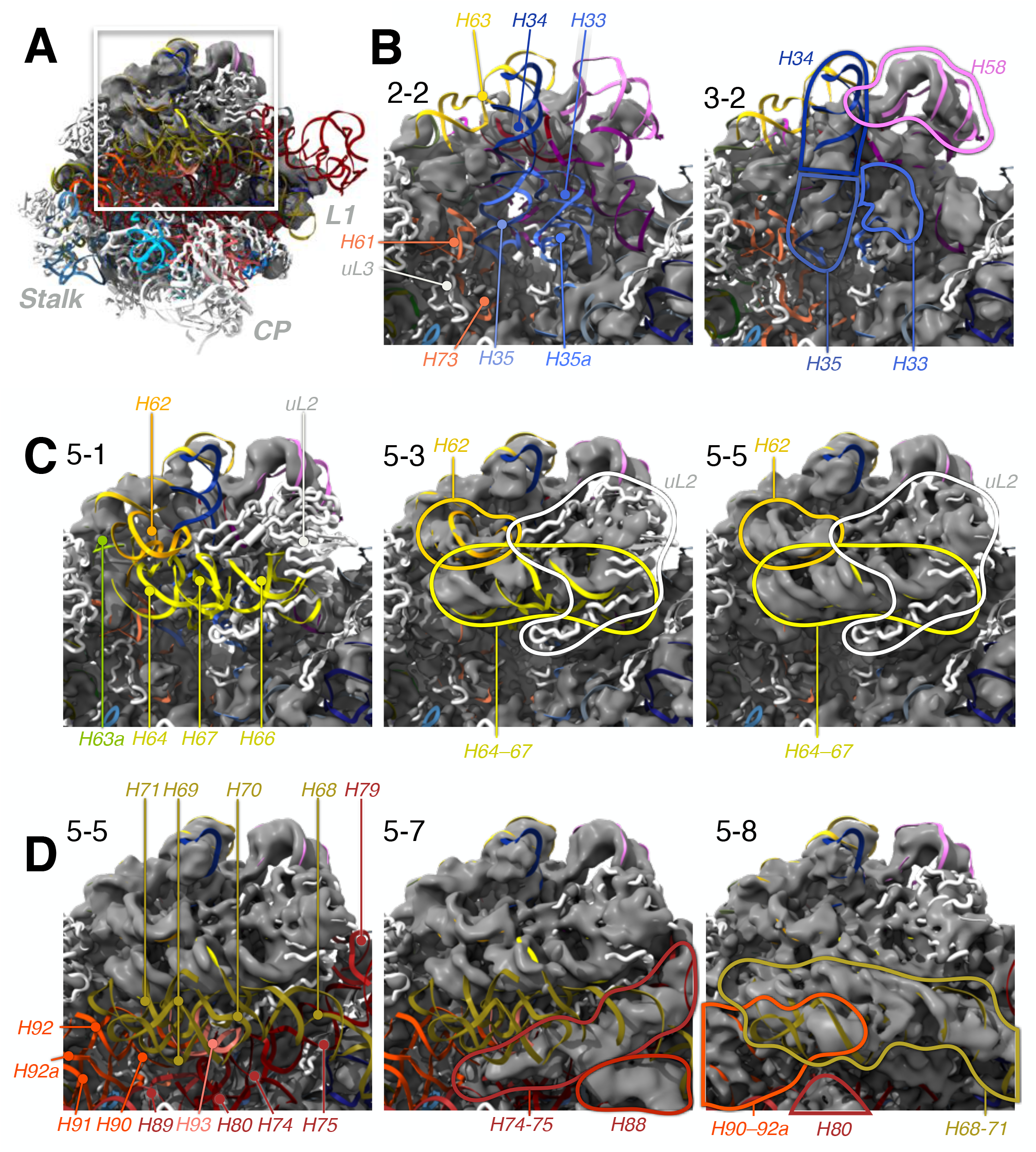
Structural dependencies between uL2 and 23S rRNA domains IV and domain II. Maps (gray) are filtered according to estimated local resolution and displayed at a threshold of 0.08. The reference model of the mature particle (PDB ID 4YBB) is colored with RNA as in Figure 2G and proteins in white. Details and additional views in Figures S16, S17. (**A**) Overview showing the relevant region in the consensus reconstruction. (**B**) Helices H33–35a (blue) and H58 (pink) are absent in class 2-2 (also see Figure S17B), but is present in class 3-2. (**C**) Folding of helices H62–67 (yellow) correlates with presence of r-protein uL2 (white). (**D**) The 3’ half of domain IV (H68–71, visible in 5-8) folds outside of H74, H75 and H88 in domain V (visible in 5-7). In 5-8, there is also density for H90-92a and H80.

#### Sub-domain III_tail_ can stabilize sub-domain IV_5’_ in absence of uL2

Class 1-1, the smallest reconstructed particle, only consists of domains I (except H25), III, VI, 0 (H26a and H61) and IV_5’_ (particularly strong density for H63) (Figures S15A, S16A). This is remarkable, since folding of sub-domain IV_5’_ in other classes coincides with presence of uL2 (see below). Class 1-2 has additional density for the core of domain II, a more mature-like domain 0 and H25 of domain I, but less density for domain IV and the distal half of domain III (helices H54–59, hereafter called sub-domain III_tail_ (26)).

Based on these structures, we hypothesize that in lieu of uL2, sub-domain III_tail_ can stabilize sub-domain IV_5’_ through contacts between H57 and H58 in domain III and H63 (Figure S15B). The same correlation between the folding of sub-domains IV_5’_ and III_tail_ was observed under bL17 depletion (11), and domain III was shown to fold independently of r-proteins or the rest of the 23S rRNA (27). Sub-domain III_tail_ is mostly folded in all of the particles, independently of domain IV and uL2, suggesting that this subdomain promotes folding of domain IV rRNA at the early stages of ribosome biogenesis.

Class 2 particles show higher stability, homogeneity and resolution than class 1, probably because additional r-proteins stabilize the RNA structure. Unlike class 1, they show no density for sub-domain IV_5’_ (Figures 6B, S16A). In class 2-1, the whole flank opposite to the CP (sub-domain III_tail_ and helices in domains 0 and VI) is shifted 20 Å away from the subunit interface compared to the native particle, whereas in class 2-2, this lobe is instead in near-native position (Figure S15C).

The exit tunnel is expanded in the smaller particles, in particular in class 2-2 (Figure 3D-F). Classes 1, 2 and 3-1 are thus possible substrates for RlmF, where the A1618 modification site is accessible. These constitute 17% of the total number of particles, in reasonable agreement with 20% of the particles being labeled after *in vitro* methylation.

Class 3-1 is rather similar to 2-1 but H63 is more pronounced and there is more diffuse density in the H33–H35a region (Figure S16A). Correlated to this, domain III_tail_ is closer to its native position, stabilized by H63 (Figure S15A) and possibly by the higher occupancy of uL3 and bL17.

In class 3-2, domain II is in a more native position compared to 3-1. It is the smallest particle with unambiguous density for H34–H35 (Figure 6B), correlated with improved positioning of nearby domain III (H52, H56 and H60) and H96 and H101 in domain VI (Figure S15).

#### R-protein uL3 is important for the stability of domains 0 and VI

Class 5-1 shares attributes with classes 2-2 and 3-2, but H63 is close-to-natively folded (Figure 16A). Similar to in larger class 5 particles, diffuse density extends from H73 across the particle to H22 (see below).

Higher occupancy of uL3 in 5-1 than in 3-2 seems to cause stronger density for domains 0 and VI (except H96). The same trend is observed between 2-2 and 2-1. In the mature particle, the 128-153 loop of r-protein uL3 directly interacts with several of these helices (Figure S13B), supporting the importance of uL3 in maintaining the native H73 structure, preventing its misfolding (see above).

#### R-protein uL2 stabilizes sub-domain IV_5’_

Class 4 has many similarities to 5-1, but the misfolded H73 dislocates a lobe at the solvent-side close to the base of the stalk, including H1, H41, uL13, bL20 and bL21. Class 4 also has significantly stronger density for sub-domain IV_5’_ and more mature-like H33–H35a (Figure S16A). The differences in domains II and IV seem correlated with increased occupancy of uL2, also seen in classes 5-2, 5-3, 5-5 and 5-6 (Figures 6B, S16B). This is also seen in the clustering analysis (Figure S12A-B, rows 85 and 89, respectively).

Class 5-2 is highly similar to class 4, but with stronger density for sub-domain IV_5’_(Figure S16).

#### Helices H79 and H88 anchor the core of domain V

Class 5-3 combines features from 3-2, 5-1 and 5-2 (Figure 5). H72 and H73 are folded (Figure S13), and on the other side of the particle, the kissing-loop contact between H88 and H22 is connected via diffuse density to H73. Similar to the nearby H29 and H31, H88 is shifted from its native position in absence of the stabilizing proteins uL15, bL33 and bL35. Out of these, association of bL33 and bL35 coincide with large structural rearrangements of the CP during *in vitro* reconstitution (17). Class 5-3 has density for r-protein uL19 and low levels for neighboring uL14 (Figures S11, S15), similar to the first particle along the C-folding pathway in (11) (Figure S9).

Class 5-4 is quite similar to class 5-3, but with slightly weaker sub-domain IV_5’_ and uL2 (Figure S16B). Domain V is on the other hand more well defined, in particular H79, H88 and the proximal end of H89. Diffuse density connects these elements, which extends towards H73.

Class 5-5 shows less distinct density for domain V compared to 5-4, although H79 seems ordered. Instead, the density for uL2 and sub-domain IV_5’_ is stronger (Figure S16B), placing it in a different path in Figure 5. The distal part of helix H67 is well-defined, but the 3’-half of domain IV is disordered.

#### Stability of sub-domain IV_3’_ correlates with stability of domain V in the largest particles

In class 5-6, all the elements that are visible in either of the smaller particles in class 5 are ordered (Figures 5, S16).

Class 5-7 shows strong and more well-defined density for helices H74 and H75 in domain V, connecting all the way from H73 to H88 and H79 (Figure 6D). H79 is packed against H10, but they are shifted from their native position in absence of bL28. H76 also has strong density, extending into the L1 protuberance formed by the folded but flexible helices H77–78 (Figure S5). Similarly, folding of domain V along the C/E paths starts with H79 to expand towards the two protuberances under bL17 depletion (11). The higher occupancy of uL13 in class 5-7 seems to stabilize H1 and H41 while, as in other classes, the relatively strong density for uL14 and uL19 does not appear to influence the rRNA.

Class 5-8 shows more native structure for domain V, which is partly held in place by sub-domain IV_3’_ (Figure 6D), but the distal end of H69 is flexible. There is also weak density for the rest of the 3’ end of domain V, including H90–93 and the proximal end of H89. Native positioning of H89, however, requires uL16 (17). Finally, the H82–H87 branch of domain V, that forms part of the CP, shows partial order, and very weak density can be discerned for additional large CP elements.

#### The CP is severely destabilized in the LiCl core particles

Class 6 is the only particle where most of the CP rRNA, including 5S rRNA, is folded, but otherwise similar to class 5-8. The CP is highly depleted of r-proteins, but contains low levels of uL5, uL18 and uL30. Relative to the mature particle, the entire CP is shifted away from the subunit-interface side.

### Folding of helices H49a and H35 demonstrates analogies between ribosome disassembly and assembly

Step-by-step comparison of the smaller to larger particles allows the folding of parts of the LSU to be modelled as an intricate sequential addition of structural layers onto a pre-existing core. As an example, we analyzed the folding of helices around H49a (modification site of RlmF in domain III) and H35 (domain II) (Figures 7, S18). These helices together form tertiary interactions between domains 0, II, III, IV and V in a region stabilized by the positively charged extensions of r-proteins uL2, uL22 and bL34.

**Figure 7.**
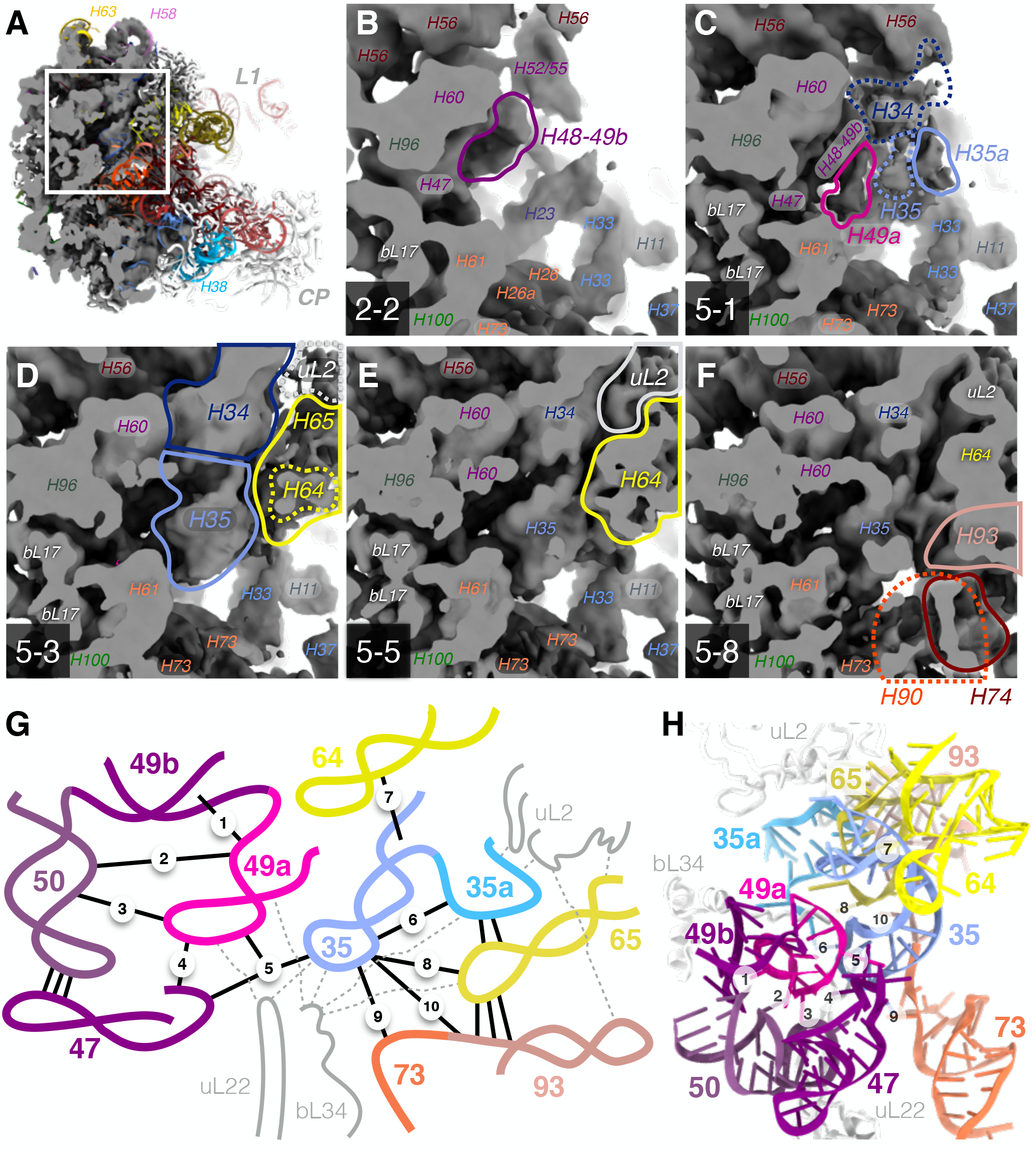
Folding of the H49a-H35 region in different particles suggests a local order of assembly. (**A-F**) Maps showing varying degrees of folded structure in this region. Lines indicate density that differ between maps (dashed line indicate weak/diffuse density). The maps are not sharpened or low-pass filtered and shown at level 0.05. Figure S18 shows the same view for all particle classes. (**A**) The overview shows the consensus map and a mature model, PDB ID 4YBB, colored as in Figure 2G. (**B**) In class 2-2, H48 is folded, but not H49a. (**C**) Density for H34, H35 and H35a outside of H49a in class 5-1. (**D-E**) Occupancy for H64, H65 and uL2 is partial in class 5-3, but close to full in 5-5. (**F**) Helices H90, H93 and H74 are folded in class 5-8. (**G**) Schematic view of stabilizing interactions in the H49a-H35 region. Tertiary contacts for H49a and H35 are indicated with solid lines and numbered in approximate assembly order, as deduced from the maps (see text for numbers). Dashed lines indicate contacts to r-proteins. (**H**) Structure of helices and r-proteins in (G) in the mature 50S (PDB ID 4YBB).

#### Folding and tertiary interactions of H49a

Helix H49a is a stem-loop in domain III that bridges to domain II, close to the nascent-chain exit tunnel (Figure 3D, F). In the mature particle, the stem of H49a packs against helices H49b and H50 and the loop forms interactions with H50, H47 and H35 (Figure 7G). Although most of domain III is folded across all classes, several different conformations and tertiary contacts of H49a are observed (Figure S18), allowing reconstruction of the sequential assembly of interactions in this region (Figure 7B-F). In class 2-2 (Figure 7B), H48 and H49b are folded but H49a is completely disordered. In other early classes (2-1 and 3-1), H49a is folded but points away from H50 at different angles. In class 5-1 (Figure 7C) and 3-2, H49a packs against H49b (contacts 1 and 2 in Figure 7G). Although the loop of H49a lacks most of the native tertiary contacts, there is weak density for A1616 connecting to H50 (contact 3). Class 4 is similar but also shows density for the 5’-end of H47 (contact 4) and for H35. In class 5-3 (Figure 7D) and 5-2, H49a is in a close-to-native conformation. Contact 5 is formed through the stacking between A1618 of H49a and A749 of H35 observed in class 5-4, and the triple-A stack is completed by A1272 of H47 as observed in class 5-5 (Figure 7E). The stack would be further stabilized by the N6 methylation of central A1618 (28). Classes 5-6 to 5-8 (Figure 7F), show only a slight distortion of the H49a loop at the flipped-out base of A1614 compared to the native structure where it forms an interaction with the extended loop of uL22, with rather weak density for the tip in all classes.

#### Folding and tertiary interactions of H35

Helix H35 is a stem-loop in domain II, which packs outside H49a, contacting domains 0 (H73), IV (H64 and H65) and V (H93) (Figure 7G & 3F). H35 is disordered in most small classes such as 2-2 (Figure 7B) and only vaguely discernable in the other small particles. The stability of H35 shows correlation to presence of r-protein uL2 (Figure 7C-E). uL2 interacts with H35a, which packs outside of H35 (Figure 7C), and in turn contacts both H35 (contact 6) and H65 (Figure 7D). An A-minor contact interlocks H64 and H35 (contact 7), observed in the more mature-like classes 5-3 and 5-5 (Figure 7D, E), but absent in *e*.*g*. class 5-1. The further interactions of H65 with the loop of H35 (contact 8) and of its loop to H73 and H93 (contacts 9 and 10) and to uL22 are only present in the largest particles such as class 5-8 (Figure 7F), completing the native structure.

## DISCUSSION

Core particles of the LSU, where the most loosely attached ribosomal proteins and the 5S RNA have been removed using high-salt wash protocols were first used in studies of ribosome assembly more than half a century ago (29, 30). We here provide structural understanding of what these particles are and investigate their similarities and differences to structures occurring during *in vivo* and *in vitro* ribosome assembly.

We have here shown that removal of loosely associated r-proteins with a high-salt wash allows access to the smallest stable cores of the large subunit. These particles show density for only 4–8 r-proteins, but additional ones may be bound to disordered rRNA regions. With exception of the smallest particles, the reconstructed cores show strong similarity to intermediates isolated from bL17-deficient cells or from *in vitro* reconstitution (Figure S9) and their protein content agrees with *in vivo* assembly (6). These similarities between states obtained during disassembly and different variants of assembly strongly support the innate propensity of the rRNA and the r-proteins to adopt distinct, meta-stable complexes. Evolutionary, this inherent property is likely important for the robustness of ribosome assembly upon changing or challenging conditions, such as disabling mutations of an r-protein (11). Still, local minima in the energy landscape can lead to trapping of misfolded structures. One such example is the strong non-native secondary structures observed to form by the 3’ strand of helix H73 (Figure 4). This manifests the need for ribosomal proteins not only to facilitate formation of the active RNA structure during co-transcriptional *in vivo* assembly, but also to maintain the native structure in a partly folded structural context. During *in vivo* assembly, RNA helicases allow remodelling of such misfolded structures (2) and during *in vitro* assembly the optimized conditions of temperature and magnesium (4, 31) may prevent their formation.

The r-protein occupancy of our LiCl core particles largely agrees with previous characterizations (5, 7, 19, 30). The spontaneously exchangeable r-proteins *in vitro* and *in vivo* (8) are the most loosely associated ones, a small subset of the ones that can be dissociated with salt. There is no indication that their dissociation and re-association would cause major structural changes, which seems evolutionary important to maintain the active pool of ribosomes.

The wide range of LSU sub-particles that we observe allows analysis of the consequences of removal of specific r-proteins. For some r-proteins, *e*.*g*. uL3, a primary binder, and uL2, a later binder (6) there is a strong link between their presence and observation of the native rRNA structure in their close surrounding. The central protuberance shows high sensitivity to deproteination and unfolds as proteins leave. In contrast, it can fold early during ribosome biogenesis as demonstrated in the bL17-depletion model (11) and during LSU *in vitro* reconstitution (17). The structural stability of domain III seems less dependent on r-proteins.

To what extent does ribosome disassembly *in vitro* reverse the process of ribosome assembly *in vivo* or *in vitro*? Apart from the CP-region proteins uL5, uL15 and uL18, the r-proteins that bind first during assembly are also the ones that stay bound during high-salt wash. The disassembly process is sterically constrained by outer assembly layers of the ribosome, and thereby perhaps more similar to *in vitro* assembly where the structure forms starting from the core and continuing outwards. During the co-transcriptional *in vivo* process there is an additional directionality, where at a given time only part of the rRNA can be available for folding and tertiary interactions (reviewed in (2)). Still, the correlation between occupancy of r-proteins in our core particles early binding during *in vivo* assembly (6) suggests that directionality is not a main factor in r-protein affinity.

The reconstructed series of particles subjected to r-protein dissociation corroborates the multi-pathway of assembly (11, 32, 33) by demonstrating the semi-independent stability of folding blocks of rRNA and r-proteins. In particular, since parts of domains IV and V can remain structured while the other unfolds, we can from the LiCl core particles derive two distinct pathways where either of these blocks independently remain folded (Figure 5).

Compared to the well-studied early, co-transcriptional 30S assembly process (34), little is known about the corresponding steps of 50S assembly. However, this study provides examples of concepts that have been described for the 30S such as the sequential reduction of structural variability of RNA by binding of r-proteins, specifically in regions that require long-range interactions to reach their native structure. There is a clear similarly between how r-proteins during assembly recognize and bind to structured regions and further contribute to the maturation of the nearby structure, and how dissociation of r-proteins seems to start with the disordering of their extensions and loops coupled to loosening of rRNA structure to be followed by dissociation of the folded r-protein domains and unfolding of the respective rRNA binding sites. The native packing of the RNA structure sometimes requires such r-protein tails or loops, as an example H49a does not adopt its fully native conformation in absence of the uL22 loop (Figure 7F), the deletion of which has been shown to lead to accumulation of immature LSU particles (35). In addition, we observe a multi-pathway disassembly process, in agreement with the complex “assembly landscape” of the 30S subunit that was proposed as analogous to the energy landscape in protein folding (36). These observations support that the core particles in some respects can be used as mimics of transient early assembly intermediates, for example in studies of maturation factors.

In conclusion, the *in vitro*-generated 50S LiCl core particles represent a collection of intrinsically stable or meta-stable complexes, from which we can learn about inherent structural properties of ribonucleoprotein particles, and specifically about states that are likely to occur during 50S ribosome assembly.

## SUPPLEMENTARY DATA

Supplementary Data are available at mBio online.

## ACKNOWLEDGEMENTS

This work was supported by grants 2016-06264 and 2017-03827 from the Swedish Research Council to MS.

The cryo-EM data was collected with the aid of Julian Conrad and Marta Carroni at the Solna node of the Cryo-EM Swedish National Facility funded by the Knut and Alice Wallenberg, Family Erling Persson and Kempe Foundations, SciLifeLab, Stockholm University and Umeå University.

The authors declare no conflicts of interest.

SK performed protein purification and prepared the sample with DL. DL prepared ribosomes and did all cryo-EM work. MS conceived the project and obtained funding. DL and MS wrote the manuscript. All authors approved the manuscript.

